# Fourier Helicity Spectra as Quantifiers of Multiscale Chirality

**DOI:** 10.64898/2025.12.15.694500

**Authors:** Sang Won Im, Jessica Ma, Neel Moudgal, Nicholas Kotov

**Affiliations:** Department of Chemical Engineering, University of Michigan; Ann Arbor, 48105, USA; Center of Complex Particle Systems (COMPASS), University of Michigan; Ann Arbor, 48105, USA; Department of Biophysics, University of Michigan; Ann Arbor, 48105, USA

## Abstract

Quantifying the chirality of three-dimensional (3D) objects is crucial for understanding interactions of nanoscale structures, biological molecules, and hierarchical materials. Although chirality is simultaneously present at multiple scales for each case, existing chirality measures rely only on singular scale-specific quantifiers, and some of them yield incorrect chirality assignments for biomolecules. Here we introduce the Fourier Helicity Spectra, a mathematical framework to decompose geometry of chemical and biological structures encompassing the scale dependence of mirror asymmetry. Fourier decomposition of 3D objects into helical harmonics enables one to compute a handedness-resolved characteristics of complex shapes, while reducing computational cost. We show its applicability for a diverse range of chemical systems, from biomacromolecular electron densities to tomographic reconstructions of inorganic nanostructures. Furthermore, inverse transform of isolated spectral components into real space enables visualization of helicity-matched interfaces, identifying how chirality governs biomolecular interactions. We anticipate that this methodology can be generalized with alternative basis sets to encompass a broad range of chiral symmetries governing biological signaling, polarization optics, drug design, and advanced synthesis.

## Main Text

Chirality, or mirror asymmetry, is a fundamental geometrical property that can be transferred via self-assembled from the angstrom scale to the macroscale, traversing along the way multiple building blocks of life. Between small molecules and large organisms, there are multiple intermediate structures exemplified by DNA, peptides, protein assemblies, organelles, and cells that display multiple levels of chirality. For instance, DNA possess mirror asymmetry both at the level of base pairs and the assembled macromolecule overall. The presence of multiple scales of chirality was also demonstrated for self-organized biomimetic nanostructures, which display signatures of angstrom-, nano-, meso-, and micro-scale chirality, as observed in both microscopy and spectroscopy.(*1*) The ability of chemical structures to be left-handed at one scale and right-handed at another scale at the same time requires quantification of chirality that should embrace its scale-dependent variability. The need for in-depth quantitative understanding of chiral relationships and their dimensional hierarchy in living and non-living matter is further amplified by the difficulties of protein folding algorithms with right-handed amino acids,(*2*) a realistic possibility of ‘mirror life’,(*3, 4*) and numerous technological needs for materials possessing chiral features with various dimensions.(*1, 5*–*12*).

However, both biology and chemistry predominantly use binary chirality classifiers, exemplified by *L*- and *D*-amino acids, sugars, and other compounds. Various mathematical concepts have been suggested to quantify chirality, such as the Hausdorff chirality measure (HCM),(*13*) Osipov-Pickup-Dunmur index (OPD),(*14*) continuous symmetry measure (CSM),(*15*) Petitjean chiral index,(*16*) graph theoretical chirality measure (GTC),(*17*) and helicity measure.(*18*) All of them consider a singular metric to be the comprehensive descriptor of mirror asymmetry of chemical and biological structures, which contradicts the large body of widely available experimental knowledge. Furthermore, HCM and CSM are enantiomer-agnostic, i.e. quantify only the magnitude of chirality without distinguishing handedness. The chiral measures that distinguish different enantiomers by their positive or negative sign, such as OPD, suffer from “chiral zeros,” a mathematical artifact when mirror asymmetric shapes return OPD = 0. This issue worsens for molecules with high molecular weight, making pseudoscalar singular value metrics inapplicable for most cases. Importantly, neither HCM nor OPD gives any practically relevant correlations with basic biological properties, such as the ability of proteins to form complexes.(*19*)

To address these fundamental problems, we introduce here the Fourier helicity spectra (FHS), a measure for multiscale chirality inspired by the Fourier transform (FT) decomposition of complex mathematical functions into constituent harmonics. To capture the chiral properties of an object, we take advantage of three-dimensional (3D) functions with helical and cylindrical shapes that, together can reproduce the complex shape of objects in the smallest details. In other words, the orthogonal expansions of the object in a cylindrical coordinate system, with rotational, axial, and radial modes serve as the basis functions for FHS. Compared to all current chirality measures, the wavelength of the basic functions provides direct connection to the scale of the object, while strict helicity of their shapes enables handedness-aware enantiomer-specific chirality measure without chiral zeros. The commonality of helices, spirals, and helicoids in Nature, as exemplified by DNA, α-helices, proteins, crystals,(*20, 21*) nanoparticles,(*1, 10*–*12*) bacteria, plants, etc, makes helical functions a logical choice for quantitative description of chirality relations in biology. We demonstrate these relationships for protein–protein and protein-DNA interactions,(*22*) which can be extended to other structurally complex and dynamically reconfigurable molecular systems.

## Results and discussion

### Orthogonal expansions with helical base-functions

The FHS for a volumetric density *ρ*(*r,φ,z*) defined in cylindrical coordinates is given by

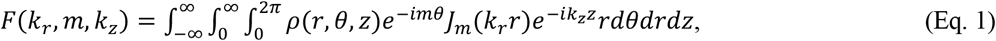

where *J*_*m*_ denotes the *m*-th order Bessel function of the first kind, *k*_*r*_ and *k*_*z*_ are the radial and axial wave numbers, and *m* denotes the azimuthal harmonic. The Bessel function naturally arises as the radial eigenfunction when solving the Helmholtz equation in cylindrical coordinates, ensuring orthogonality and completeness of the basis set under axial symmetry (**Fig.1a**). This representation preserves explicit definitions of pitch and handedness through the relationship between *m* and *k*_*z*_ with pitch defined as *p* = 2π*m*/*k*_*z*_ Modes with positive pitch (*mk*_*z*_ > 0) are left-handed, whereas modes with negative pitch (*mk*_*z*_ < 0) are right-handed (**Fig.1b**). In our implementation, scalar fields or voxel data, typically defined in Cartesian coordinates, are first converted into cylindrical coordinates resulting in the volumetric density function *ρ*(*r,φ,z*). Then, we apply one-dimensional (1D) fast Fourier transforms (FFTs) along the *φ* and *z* axes and a quasi-discrete Hankel transform (QDHT) (*23*) along the *r* axis. The QDHT enforces Parseval energy conservation by exploiting the orthogonality of Bessel functions at nodes tied to specific Bessel zeros. Note that a precomputed transform matrix constructed from those zeros results in matrix–vector multiplication with *O*(*N*_*r*_ *N*_*φ*_ ^2^*N*_*z*_ ) scaling, where *N* _*r*_, *N*_*φ*_, and *N*_*z*_, are the numbers of samples along each axis. Consequently, the computational time for FHS scales as *O*(*N*^4/3^) making its calculation faster than those for HSM, OPD, CSM and other chirality measures for equal number of voxels.

**Fig. 1.**
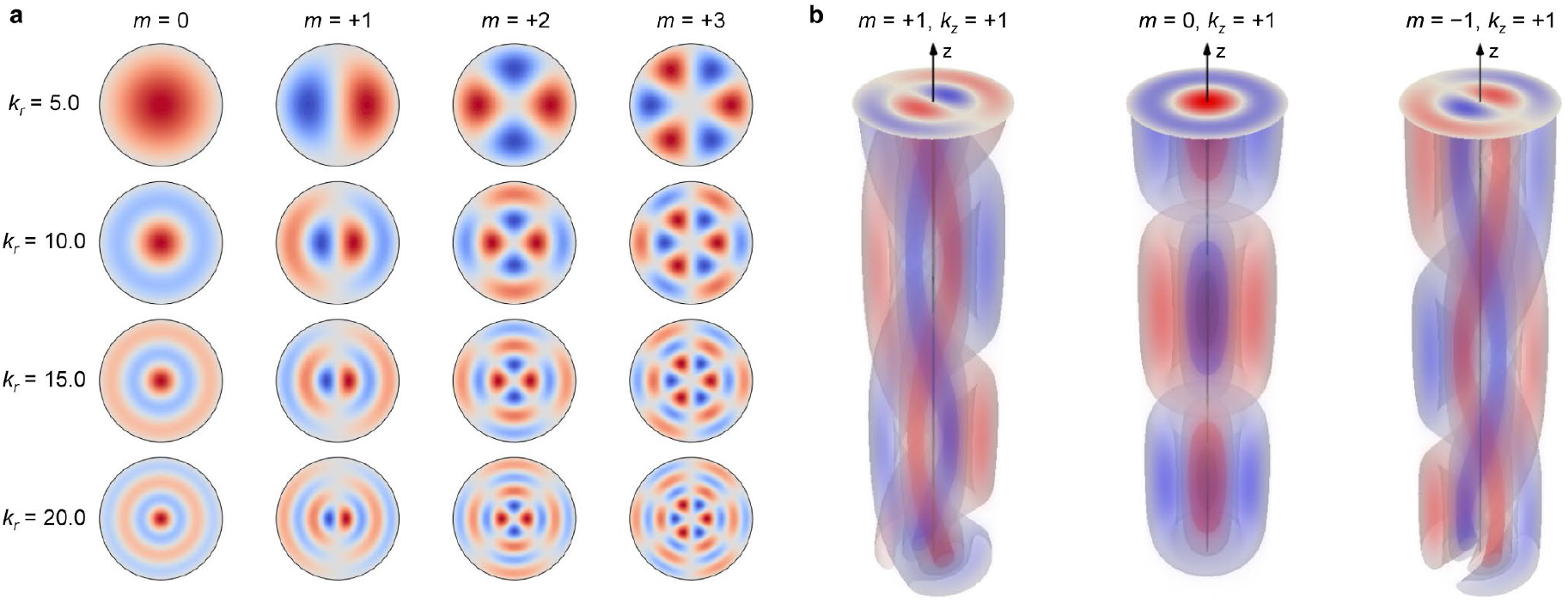
Base functions for the Fourier helicity spectra. (**a**) z = 0 slice of the cylindrical Bessel function of the first kind *J*_*m*_(*k*_*r*_*r*) with azimuthal order *m* and radial wavenumber *k*_*r*_. (**b**) Illustration of helical basis functions *J*_*m*_(*k*_*r*_*r*)exp(*i*(*mϕ+k*_*z*_*z*)) with different chirality: left-handed (*mk*_*z*_ > 0), achiral (*mk*_*z*_ = 0), and right-handed (*mk*_*z*_ < 0). *k*_*r*_ is the second zero of the first-order Bessel function.

Unlike other chirality measures, FHS resolves 3D geometries into modes indexed by (*k*_*r*_, *m, k*_*z*_). Among these parameters, *k*_*r*_ sets the radial oscillations encoding radius and thickness. The coupling of *m* and *k*_*z*_ defines handedness and pitch of helical mode. For a helix with fixed radius *R*_0_ and pitch *p = P*_0_, the periodicity along the *z* axis coupled with the azimuthal rotation leads to discrete modes in *m*-*k*_*z*_ spectrum that satisfy the relationship *k*_*z*_=2π*m*/*P*_*0*_, each mode corresponding to an integer *m* (**Fig.2a**). These modes appear as ‘layer lines’ in *k*_*r*_-*k*_*z*_ spectra, where each has radial intensity of |*J*_*m*_(*k*_*r*_*R*_0_)|^2^ peaking at *k*_*z*_ = ±(2π*R*_0_/*P*_0_)*k*_*r*_. Note that the X-shaped profile of these spectra is directly related to the X-ray diffraction patterns observed for DNA fibers that catalyzed the discovery of the helical shape of this foundational molecule.(*24, 25*)

**Fig. 2.**
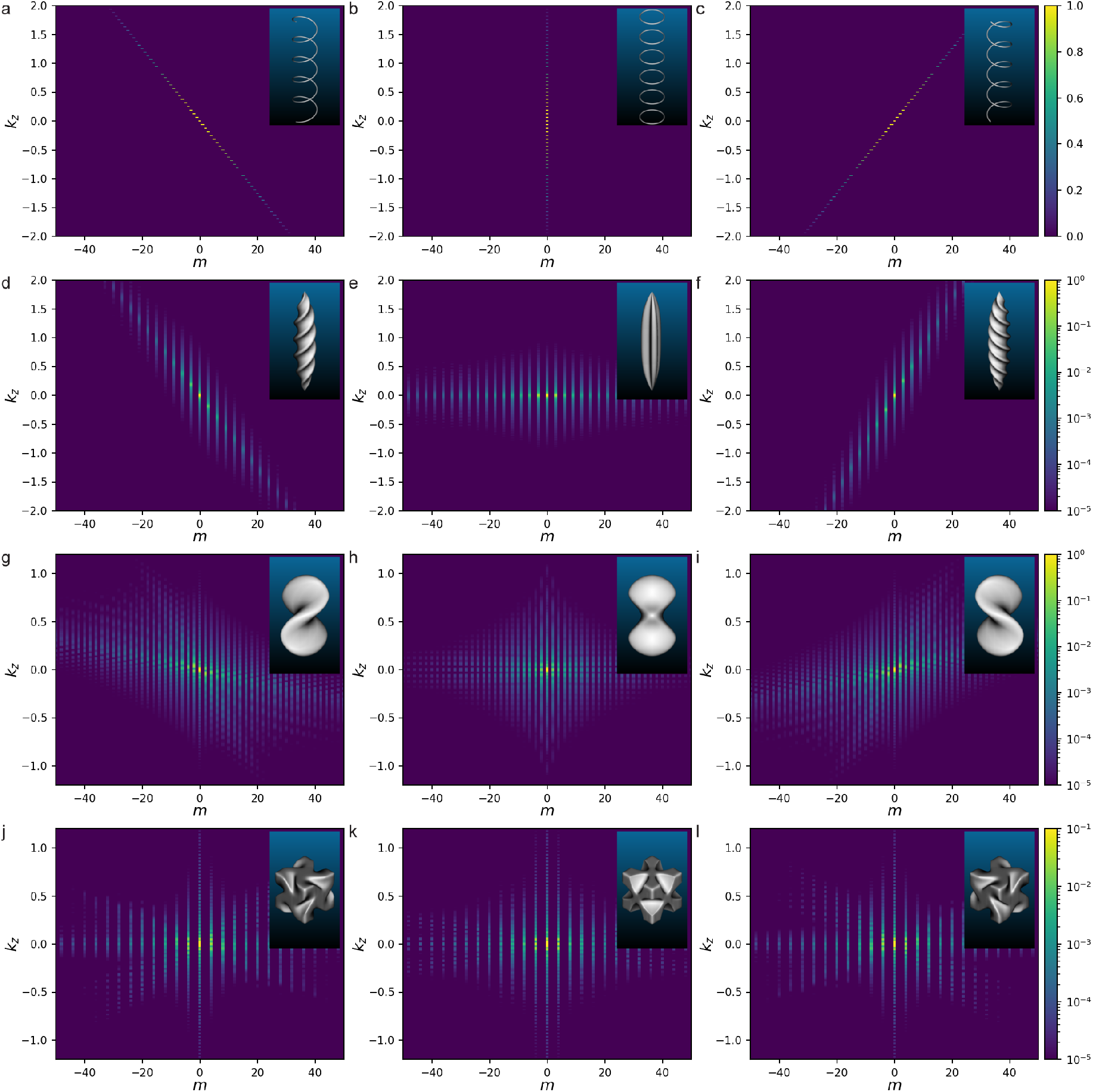
Fourier helicity spectrum analysis of 3D chiral and achiral geometries. *m-k*_*z*_ spectrum for right-handed, achiral, and left-handed (**a-c**) helices, (**d-f**) tapered helices, (**g-i**) bowtie particles, and (**j-l**) 3D helicoids are shown. 3D renderings of the corresponding structures are included as insets.

The *m-k*_*z*_ spectra describe helical modes quantitatively and efficiently as the slope of linear ridge directly captures the pitch through the relation *m*/*k*_*z*_ = *p*/2π, while the sign of the slope captures the handedness. Helices with different pitches generate lines with different slopes. The mirror reflection of the helix that generates an opposite enantiomer reverses the sign of the slope. By contrast, a stack of achiral rings with axial spacing equal to the pitch preserves the same layer-line spacings in *k*_*z*_ but concentrates the intensity along the *m* = 0 line (**Fig. 2b,c**).

Beyond ideal helices, FHS was applied to complex 3D chiral objects to resolve their helical motifs. For example, we calculated the FHS of a tapered helix with three helical grooves, which is a typical morphology observed in assemblies of tetrahedral nanoparticles.(*26, 27*) The Bessel function of expansion their enantiomers results in linear *m-k*_*z*_ spectra centered at *k*_*z*_=2π*m*/*p*, where the slope is determined by the pitch as in the ideal helix (**Fig. 2d**). The need for greater number of helical harmonics to produce the non-constant pitch in tapered helices than in ideal infinite helices, manifests as broadening of the vertical features along the *k*_*z*_ axis by ∼2π/*L*, where *L* is the length of the finite tapered helix. Simultaneously, the constrain on *L* induces a spread of slopes, producing a fan-shaped pattern. Importantly, the mirror symmetry of the solid core of the tapered helix that is nearly cylindrical in shape contributes to achiral modes, adding intense components at *m* = 0 and *k*_*z*_ ≈ 0. The presence of *n*-fold rotational symmetry results in the attenuation of azimuthal orders that are not multiples of *n*. For example, for triple-groove helix derived from a 3-fold symmetrical cylinder, modes with *m* ≠ 3ℓ (where ℓ is an integer) are attenuated. These spectral broadening and azimuthal selection rules are equally applicable to an achiral 3-fold symmetric tapered rod, producing a horizontal line centered at *k*_*z*_ = 0 (**Fig. 2e**).

Then, we applied FHS to the even more complex geometries of experimentally observed nanoscale and microscale particles with shapes of helicoids. One of them are bowtie particles observed for assemblies of twisted nanoribbons composed of cadmium ion and homochiral dipeptide cystine. They have an overall helical shape, but the pitch varies with axial position—*P*_0_ decreases near the center and increases toward the periphery. In addition, the small number of turns and the solid core distinguishes bowties from an ideal helix. In the *m–k*_*z*_ spectrum these features appear as a fan-shaped distribution of harmonics spanning a wide pitch range, with a strong achiral mode at *m* = 0 and near *k*_*z*_ = 0 due to the solid asymmetric core and pronounced broadening along *k*_*z*_ due to variation of *p* (**Fig. 2g**). Their helical expansion also indicates the presence of two dominant *p* of the same handedness, visualized as two diagonal traces with different 2π*m*/*p* tangents.

Another type of particles are the helicoids observed in the process of chiral crystal growth of gold in the presence of chiral thiol molecules such as glutathione (**Fig. 2j**). One of the special attributes of these particles compared to cases considered above, is that they do not have a distinct axial shape and are relatively isotropic as many proteins and other globular biomolecules. Their morphology can be described as 432-symmetric arrangement of tilted concave gaps along the cube edges, arising from mirror-symmetry breaking of the 4/*m* 3 2/*m*-symmetric morphology of the gold crystal.(*10, 28*) With [001] chosen as the axis, the four parallel edges correspond to positive pitch, whereas the eight perpendicular edges correspond to negative pitch. Consequently, the *m*–*k*_*z*_ spectrum after FHS shows coexisting ridges with positive and negative slopes, together with a strong achiral mode originating from the central part of the particles (**Fig. 2j**). Similarly to bowties, their *m*–*k*_*z*_ spectrum clearly demonstrates the presence of two preferential pitches visible by the diagonal traces. Unlike bowties, the handedness of the two pitches is opposite to each other.

### Objects with multiple scales of chirality

In aggregate, **Fig. 2** indicates that *m*-*k*_*z*_ spectra can separate and quantify helical and achiral components within complex 3D geometries. It also shows that different helical harmonics with multiple pitches can be extracted from complex shapes. Going further, we need to consider that biological and biomimetic structures, can have chiral components organized hierarchically as it is typical in biology with signature chiral features at each level of organization. Such structures are often comprised of chiral components across multiple scales with mixed handedness and exhibit achiral dilution arising from nonideal geometries. Orthogonal expansion with helical basis functions with different pitch makes possible to extract scale-dependent chirality metrics specific to hierarchical geometries capturing the relative contribution of left-handed, right-handed, negative, and achiral harmonics. One can characterize the modes observed in *m*-*k*_*z*_ spectra by their energy *E*(*m,k*_*z*_) obtained by integrating |*F*(*k*_*r*_,*m,k*_*z*_)|^2^ over *k*_*r*_, which will lead to analogs of density of states in quantum systems. Furthermore, one can carry out this analysis separating |*F*(*k*_*r*_,*m,k*_*z*_)|^2^ for positive and negative pitches. Subsequently, a continuous pitch–energy density, i.e. *ρ*(*p*), spectra are computed via kernel density estimation, using per-mode bandwidths that account for the non-uniform sampling of pitch *p* induced by the uniform *k*_*z*_ grid through the mapping, which is the result of consideration of vastly different scales at the same time.

We calculated *ρ*(*p*) spectra of 3D chiral geometries for representative biological macromolecules and chiral nanoparticles (**Fig. 3)**. The 3D structures from the protein data bank (PDB) structures resolved by X-ray diffraction, or cryo-EM were used as starting geometries of biomacromolecules, followed by extraction of helical harmonics, *m*-*k*_*z*_ and *ρ*(*p*) spectra. 3D TEM reconstructions were employed for nanoparticles.

**Fig. 3.**
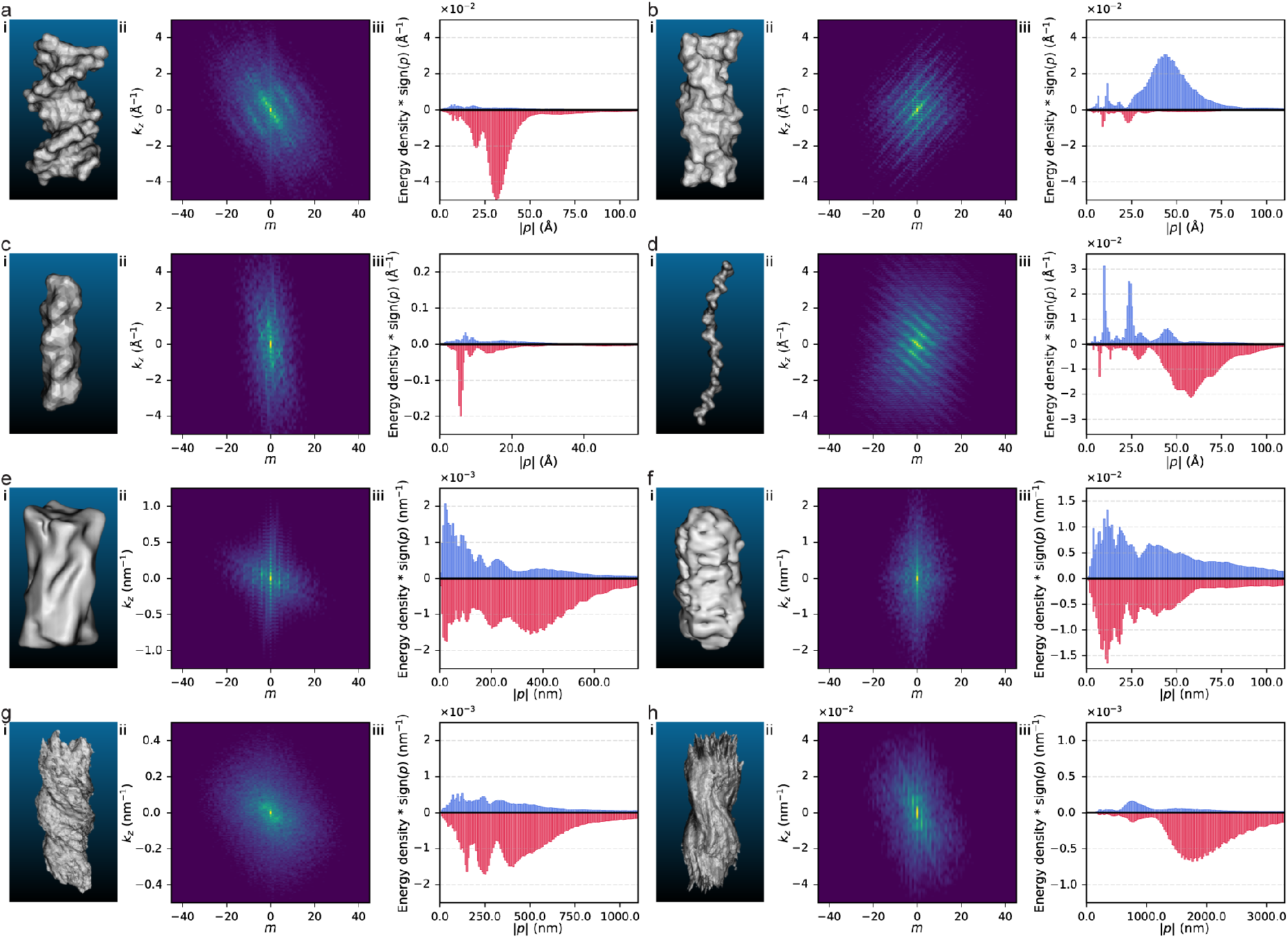
Fourier helicity spectrum analysis of various chiral nano- and microstructures, from biomacromolecules to inorganic nanoparticles. (**a**) B-DNA (PDB: 1D28), (**b**) Z-DNA (PDB: 4OCB), (**c**) α-helix (acetylcholine receptor M2) (PDB: 1CEK), (**d**) type II collagen (PDB: 1K6F), (**e**) chiral gold nanorod synthesized using L-cysteine, (**f**) micelle-directed wrinkled gold nanorod synthesized using (S)-(-)-1,1′-binaphthyl-2,2′-diamine, (**g**) tapered helical assembly of cadmium telluride nanoparticles with L-cysteine, (**h**) bowtie-shaped assembly of cadmium-L-cysteine nanoribbons. Solvent-accessible surface areas were used for the biomolecules, and transmission electron microscopy tomographic reconstructions were used for nanoparticles and assemblies. Rendered image (i), *m*-*k*_*z*_ spectrum (ii), and pitch-energy density spectrum (iii) of each structure are presented.

DNA is one of the most representative examples of helical structure of biomacromolecules. Its characteristic double helix structure is resolved as clear diagonal lines in the *m*-*k*_*z*_ spectrum and as a single peak in the *ρ*(*p*) plot. A right-handed pitch peak at 28 Å for A-DNA (*29*) (Fig. S2), a right-handed pitch peak at 33 Å for B-DNA (*30*) (**Fig. 3a**), and a left-handed pitch peak at 45 Å for Z-DNA (*31*) (**Fig. 3b**) were resolved, each matching their X-ray data. Helical secondary and tertiary structures of proteins could also be resolved by FHS. Acetylcholine receptor (AChR) M2,(*32*) an example of an α-helix, the most common secondary structure with helical geometry, yields a characteristic pitch of 5.4 Å together with additional peaks dependent on the amino acid side chains (**Fig. 3c**). Furthermore, multiscale helicity of secondary and tertiary structures of collagen (*33*) could be resolved by FHS; ∼10 Å pitch and ∼28 Å pitch for left-handed 10/3 helical secondary structure and ∼60 Å pitch for right-handed superhelix were identified (**Fig. 3d**).

For the chiral nanoparticles and their assemblies, single-particle electron tomography volume densities or surface reconstructions reported in previous publications were used.(*11, 28, 34*–*36*) The 422 helicoid gold nanorod (**Fig. 3e**), an example of uniaxial chiral geometry expressed on inorganic nanoparticle, is synthesized via secondary growth of gold nanorods of 4/m 2/m 2/m symmetry with presence of homochiral L-cysteine, yielding a 422-symmetric chiral edges and grooves. For ∼100 nm-sized nanorods, the oblique chiral edges along the length-orientation were characterized as a broad peak at right-handed pitch of 400 nm by FHS. Another example is wrinkled nanorods synthesized using the chiral-templating effect of micelle (**Fig. 3f**), which exhibit complex architectures with alternating wrinkle handedness and angles. The FHS resolved the distribution of pitches, with values from 30 to 100 nm showing higher energy in the right-handed modes.

The *m*-*k*_*z*_ and *ρ*(*p*) spectra quantifies the contributions of helical modes for a 3D structure. To quantify the excess of helical modes with specific handedness, we define a metric, the helical dissymmetry factor, that expresses the dominance of one handedness as a single value. We define the *Fourier spectral helicity* (FSH) of an overall density as a weighted sum of left- and right-handed modes, with weights corresponding to the relative energy stored in each mode

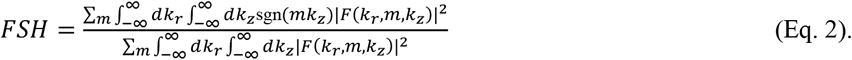

For 3D geometries, FSH lies between -1 and 1, while the ideal right-handed helix and left-handed helix with infinite length and Bessel-like radial intensity profile have FSH = −1 and FSH = +1, respectively. This metric can be computed directly from the FT coefficients with negligible computational cost. FSH can also be compared to other chirality measures. The degree of chirality of a helix is determined by the ratio of radius to pitch; when radius ≫ *p* it approaches a cylinder, when radius ≪ *p* it approaches a line, and a maximum chirality is achieved at an intermediate ratio. To reflect these properties, a definition based on torsion of helices is adopted. A helix with given pitch *p* and radius *r* has a constant torsion

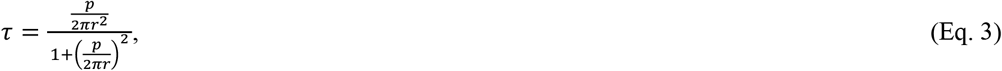

which reaches maximum or minimum value of ±1/2*r* when *p* = ± 2π*r*. We define the maximally chiral helix as one with the maximal torsion value for a given radius, which occurs when the p = 2π*r*. To achieve a unitless quantity that scales between -1 and 1, we define the *Fourier spectral chirality* (FSC) as

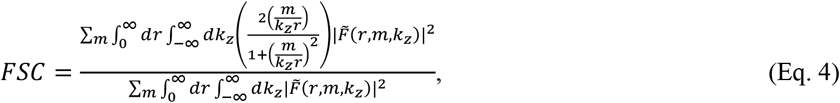

with weight function multiplied 2*r* to torsion. FSC provides a singular chirality attribute consistent with existing chirality measures and suppresses artifacts arising from helical modes approaching achiral limits.

Table 1. lists the FSH and the FSC for the examined structures. This comparison highlights the behavior of both metrics: first, as expected, structures with well-defined helical features, such as DNA, collagen, bowtie, and tapered helix, exhibit high dissymmetry and chirality. In contrast, structures with coexisting left- and right-handed contributions, such as wrinkled nanorods, or with dominant achiral components, exhibit low dissymmetry factors and reduced chirality measures. G-actin, while displaying higher HCM compared to collagen, has relatively low FSC because of its globular shape without a clear helical axis. Second, the dissymmetry factor and the chirality measures are similar for many cases. Noticeable differences arise when a specific helical mode with deviation from *p* = 2π*r* dominates the overall structure, such as collagen with elongated helical structure.

The Hausdorff chirality measure (HCM) was calculated for the geometries for the comparison. The HCM quantifies the distance between an object and its mirror by taking a minimized Hausdorff distance

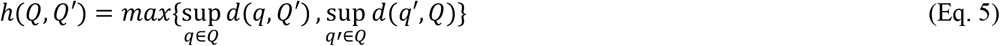

between the set of points *Q* and *Q*’ representing them. This construction returns zero if and only if the object is achiral, i.e., exactly superimposable on its mirror by a proper rotation and translation.(*13*) The absolute value of the helical chirality measure and the HCM generally show similar trends, with some deviations depending on geometries. For geometries close to an ideal helix, such as collagen or a tapered helix, the HCM is often underestimated relative to the helical chirality measure. In contrast, for 432 helicoids or certain proteins with multiple rotational axes or without specific axes, chirality value is larger for the HCM than the helical chirality measure.

**Table 1.** Fourier spectral helicity and Fourier spectral chirality of chiral structures. The Hausdorff chirality measure is provided as a comparison.

| Category | Structure | Fourier spectral helicity | Fourier spectral chirality | Hausdorff chirality measure |
| --- | --- | --- | --- | --- |
| DNA | A-form (116D) | -0.2714 | -0.2289 | 0.1606 |
|  | B-form (1D28) | -0.2785 | -0.2671 | 0.1120 |
|  | Z-form (4OCB) | +0.1942 | +0.1918 | 0.0722 |
| Proteins | Alpha helix (1CEK) | -0.0503 | -0.0434 | 0.0508 |
|  | Collagen (1K6F) | -0.2092 | -0.1061 | 0.0775 |
|  | G-actin (3HBT) | +0.0241 | +0.0244 | 0.1012 |
| Nanoparticles | 422 helicoid gold nanorod (L-Cys) | -0.0486 | -0.0487 | 0.0586 |
|  | 432 helicoid gold nanoparticle (L-GSH) | +0.0036 | +0.0010 | 0.0832 |
|  | Micelle-directed gold nanorod (S-BINAMINE) | +0.0015 | +0.0024 | 0.0531 |
| Nanoparticle assemblies | Tapered helix (CdTe, L-Cys) | -0.1340 | -0.1159 | 0.0636 |
|  | Bowtie (Cd-L-Cys) | +0.0405 | +0.0394 | 0.0655 |

### Application of Fourier helicity spectrum to complex geometries

The helical modes isolated by FHS in reciprocal space can be reconstructed on real space by inverse Fourier transform (IFT). This enables direct visualization of the 3D geometry contributed by specific modes, which is a unique feature of FHS. For example, one can visualize how left-handed or right-handed modes contribute to different regions of a 3D geometry. Mapping the difference between left- and right-handed modes of B-DNA in real space volume by IFT shows that the main double helix is dominated by right-handed (*mk*_*z*_ < 0) contributions, whereas discrete peripheral phosphate groups are dominated by left-handed (*mk*_*z*_ > 0) contributions (**Fig. 4a**). Moreover, isolating the modes by handedness and reconstructing them as 3D volumes partitions the structure into a right-handed central helix, an achiral central rod, and a left-handed outer shell (**Fig. 4b**).

**Fig. 4.**
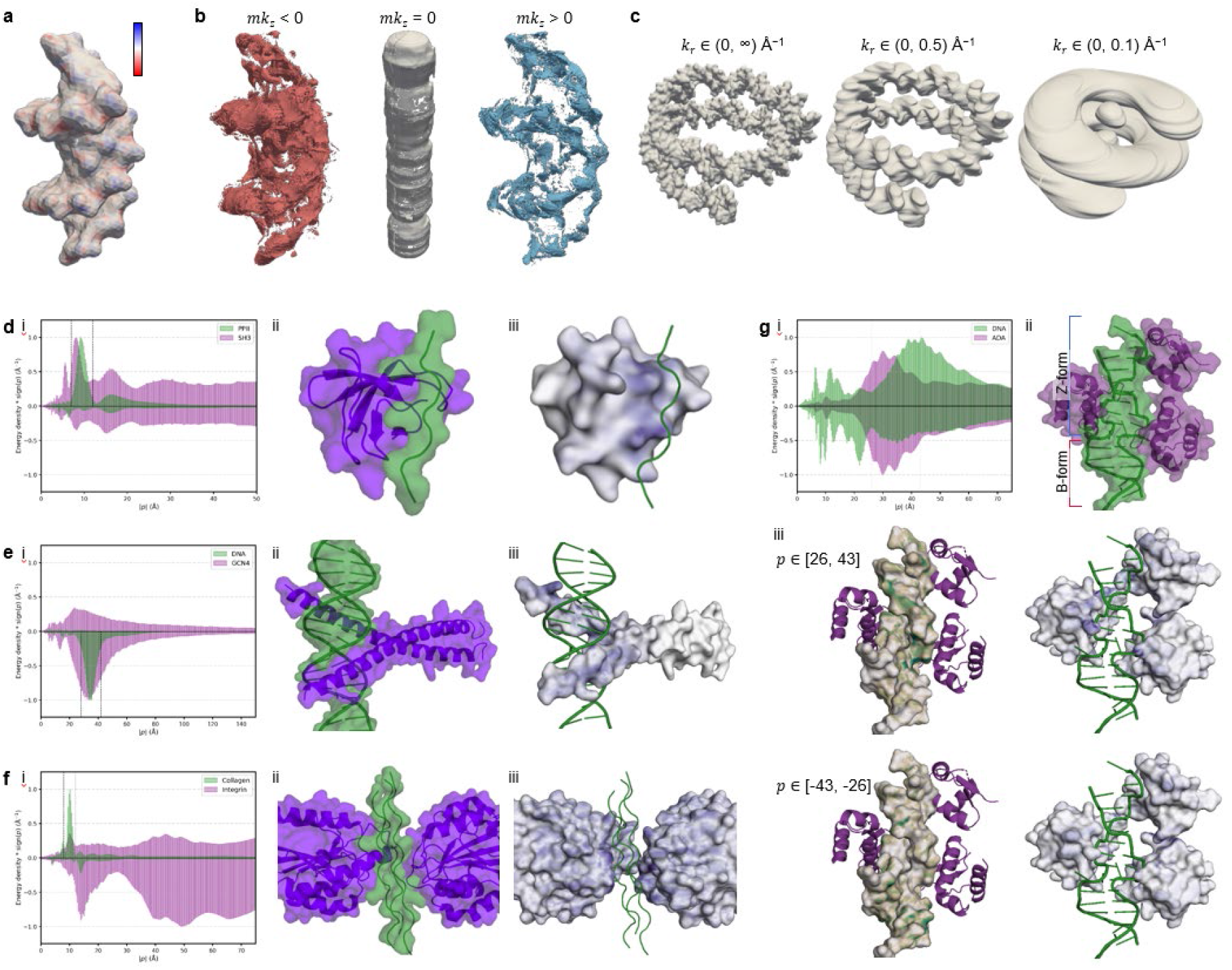
Identification of chiral components of complex geometries by Fourier helicity spectrum and inverse Fourier transform (IFT). (**a**) Mapping of helicity on the geometry of B-DNA (PDB: 1D28). The contribution of energy between left- and right-handed modes was compared after IFT onto real space. (**b**) Reconstructed achiral, left-, and right-handed modes of B-DNA. (**c**) Reconstructed geometry of wrapped DNA double helix in nucleosome (PDB: 1KX5) after applying a low-pass filter for kr. (**d**-**f**) Identification of chirality-matched lock-and-key interaction sites of (**d**) SRC homology domain and a left-handed polyproline II helix (PDB: 1SSH), (**e**) B-DNA and leucine zipper domain of yeast GCN4 (PDB: 1DGC), (**f**) integrin and the GFOGER motif of collagen triple helix (PDB: 4BJ3). For each case, a pitch-energy density plot was evaluated to identify the range of pitches matching the helical key (i). Based on the 3D structure of two compartments interacting (ii), the helical modes corresponding to the selected range of pitch were filtered and mapped onto the lock compartment using a colormap (iii).

FHS also enables multiscale analysis by simple spectral masks in radial frequency *k*_*r*_, which quantifies radial spatial variation and therefore the width of helical features. Low-pass or band-pass filtering in *k*_*r*_ followed by reconstruction by IFT enables rapid isolation of geometry with specific radial scales. This method is useful for complex chiral geometries with mixed handedness at different scales. For example, the DNA in a nucleosome is wrapped around a disk-like protein core, thereby adding an additional super-helices component with the conventional chirality of the double-helix. Because this the chirality a corresponding to component helical harmonics with larger radial range than the DNA backbone double-helix, they can be separated by applying a low-pass filter in *k*_*r*_. Specifically, a cutoff of *k*_*r*_ < 0.1 Å isolates the large-scale component that can be independently reconstructed by IFT **(Fig. 4c)**.

FHS can also be applied to a biological complex, in which interfacial recognition is governed by geometric complementarity. A representative lock-and-key complex is the binding of the Src homology 3 (SH3) domain to a left-handed polyproline II (PPII).(*37*) The pitch-energy density distributions of PPII and SH3 were analyzed (**Fig. 4d**), and the real-space amplitude of modes with pitches of ∼7-12 Å were reconstructed by IFT. As a result, the corresponding helical mode was localized to protrusions at the SH3 binding site that are complementary to the grooves of the PPII helix, indicating that matching helicity is crucial for complex formation (**Fig. 4e**). Another example is the binding of the leucine zipper dimer of the yeast transcription factor GCN4 to the helical groove of B-form DNA.(*38, 39*) When the right-handed pitch of 33 Å that corresponds to the dominant helical mode of DNA (**Fig. 4f**) was mapped on GCN4, the two N-terminal basic regions of GCN4 binding to the DNA were highlighted (**Fig. 4g**). The interaction between integrins and the triple-helical collagen provides another example of helical recognition, in which the metal ion-dependent adhesion site of integrin forms complementary conformation of GFOGER motif.(*40, 41*) Similarly, the binding sites of integrin can be identified by IFT of left-handed ∼10 Å pitch modes matching to the collagen triple helix (**Fig. 4h, i**).

Quantitative understanding of the left- and right-handed modes will also be important for complexes of nucleic acids with proteins because this is one of the most rapidly developing area of structural biology. The Z-DNA-binding domain of human double-stranded RNA adenosine deaminase I (hZα_ADAR1_) selectively binds Z-DNA at the junction with B-DNA, where Z-DNA has a left-handed (positive pitch) helix and B-DNA has a right-handed (negative pitch) helix (**Fig. 4g**).(*42, 43*) IFT of each mode with different chirality maps onto the respective parts of the DNA where the Z- and B-form segments are located. Furthermore, mapping each mode onto hZα_ADAR1_ elucidates the associated chiral selectivity and interaction, such that the left-handed mode visualizes the DNA binding site, whereas the right-handed mode maps to unrelated exterior regions. These demonstrations indicate that FHS can be a framework for elucidating chiral interactions between biomolecules. These demonstrations prove that FHS can be a framework for elucidating chiral relations governing biological interactions.

### Advantages and disadvantages of Fourier helicity spectra as measures of chirality

Compared to prior chirality and helicity measures, FHS offers the following advantages for quantification of mirror asymmetry. *First*, FHS is grounded in the concise mathematical definition of the helical basis function. This mathematical clarity yields interpretable and predictive measures with well-defined ideal geometry for maximum chirality; the geometry achieving the maximum is often not well defined for other chirality measures. *Second*, performing operations in reciprocal space via the FT introduces practical differences in robustness compared to real-space methods. Unstructured noise and high-frequency artifacts from sampling irregularities can be easily excluded. In addition, modal energies are rotation invariant, which removes errors induced by in-plane orientation jitter. *Third*, FHS offers practical computational benefits, since separable FFT and quasi-discrete Fourier methods reduce the asymptotic cost below quadratic in data size. In contrast, real-space chirality measures often require high-order pose searches or pairwise distance evaluations, for example worst-case *O*(*n*^6+*δ*^) scaling for HCM.

FHS also has disadvantages arising from the limited representativeness of approximating chirality by helicity. For example, chiral geometries with multiple symmetry axes, can exhibit chirality that depends on the definition of the axis or in which multiple modes may cancel each other. In such cases, results for multiple axes need to be combined, or the axis needs to be defined based on crystallographic orientation. The generalization of FHS to lock-and-key interactions of proteins may also be non-trivial, because the current method requires selection of a defined axis and considers only surface geometry.

## Conclusion

To summarize, FHS framework is an efficient and versatile chirality measure based on an explicit definition of the helix and computation in Fourier space. We have demonstrated FHS for diverse chiral geometries of biomacromolecules and inorganic particles, successfully quantifying their helicity and chirality. Furthermore, FHS allows the separation of helical components and the visualization of chiral lock-and-key interactions by inverse transform. We envision that this approach is not limited to geometry but is equally applicable to various scalar fields such as charge or hydrophobicity, and thus is expected to play an important role in elucidating chiral-specific interactions of proteins, molecules, and nanomaterials. In addition, our computationally efficient pipeline can be readily integrated with machine learning/artificial intelligence approaches for the development of models that explicitly incorporate chirality.

## Methods

### Fourier helicity spectrum

In cylindrical coordinates, any *L*^2^ function that is regular at the *z*-axis can be expanded in terms of the Fourier-Bessel basis

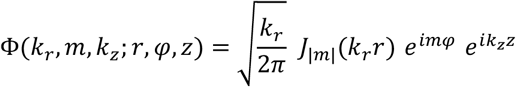

with *m* ∈ ℤ, *k*_*r*_ ∈ ℝ_>0_ and *k*_*z*_ ∈ ℝ. Here *J*_|*m*|_ is the Bessel-function of the first kind of order |*m*|. The Bessel function naturally arises as the radial eigenfunction when solving the Helmholtz equation in cylindrical coordinates, ensuring orthogonality and completeness of the basis under axial symmetry. These functions satisfy the generalized (Dirac) orthornomality condition

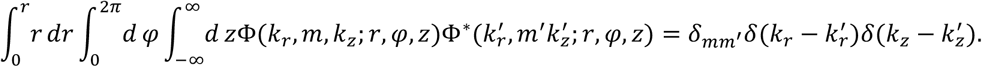

It follows that any regular *L*^2^ function *f* on ℝ^3^ can be written as

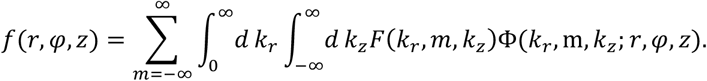

Here, we generally take *f* to be be the physical volumetric density function *ρ* of the object at hand and the corresopnding Fourier coefficients are given by:

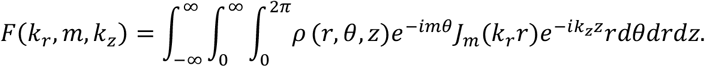

### Helicity and chirality measures

The handedness of each Fourier-Bessel mode Φ_*m*_ (*k*_*r*_, *k*_*z*_; *r, φ, z*) is encoded in sgn(*mk*_*z*_) with modes having sgn(*mk*_*z*_) = −1 being right-handed and those having sgn(*mk*_*z*_) = 1 being left-handed. Consequently, we can define the helicity of an overall density by a weighted sum of left- and right-handed modes with the weights corresponding to the relative energy stored in that mode:

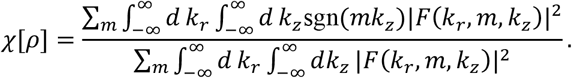

It is immediate from this definition that *χ* ∈ [−1,1] with the extreme cases *χ* = −1 and *χ* = 1 corresponding to ideal right- and left-handed helices respectively. Most realistic structures have an intermediate value of *χ* as a result of having both left- and right-handed morphological features. *χ* is thus best thought of as a weighted average of all the helical features in the structure. Structures with *χ* < 0 are predominantly right-handed, those with *χ* > 0 and predominantly left-handed, and those with *χ* = 0 have an equal amount of left- and right-handed features and thus are expected to behave as achiral. In practice, we discretely sample *k*_*r*_ and *k*_*z*_ so the integrals collapse to sums in the expression for helicity.

### Implementation details

FHS was implemented as a Python project using the FFT of SciPy and the QDHT of PyHank. FHS supports 3D geometries represented as voxels, surface meshes, or density volume reconstructed from X-ray crystallography, electron tomography, or other 3D characterization methods as well as those created by computer-assisted design software. In a typical workflow, the geometry is first aligned to the helical axis and center either manually or using the principal component analysis embedded in FHS, after which the extent and resolution of the cylindrical coordinate system are defined. The real domain extent sets the resolution in the frequency domain, and the real domain resolution sets the extent in the frequency domain. The azimuthal range is always −π to π, and the angular-frequency resolution is quantized with Δ*m* = 1. Maximum resolution can be achieved when sampling toward *r* and *z* match the resolution of raw data, and the azimuthal increment satisfies Δ*φ* ≤ Δ*r*/*L*_*r*_, where *L*_*r*_ is radial extent.

### Preparation of 3D geometric data

Helices and ideal chiral geometries were prepared by converting NURBS objects created in Rhinoceros 3D (version 7.0, Robert McNeel & Associates, Seattle, WA) with the Grasshopper add-on into meshes. For 3D geometry of nanoparticles and assemblies, the electron microscopy tomography data from previous report were used in the format as 3D density volumes or surface meshes. Surface meshes of the solvent-accessible surface area (SASA) of protein and DNA structures were obtained from the PDB using their accession codes.

### Calculation of pitch-energy density distribution

The pitch-energy density distribution plot was constructed based on the Fourier coefficients by mapping each mode (*m, k*_*z*_) to its corresponding pitch *p* = 2π*m*/*k*_*z*_. Because the transformation is nonlinear in *k*_*z*_, pitch inherits a non-uniform discretization from the uniform grid in *k*_*z*_. A first-order differential change of variables shows that *k*_*z*_ sampling interval maps to Δ*p ≈* |*dp*/*dk*_*z*_|Δ*k*_*z*_ *=* (|2π*m*|/|*k*_*z*_|^2^)Δ*k*_*z*_, which indicates coarser p spacing at low |*k*_*z*_|. For consistent energy density mapping across all modes, we estimate a continuous pitch distribution by kernel density estimation (KDE), using a per-sample Gaussian kernel width based on Δ*p*. The effective pitch resolution is controlled by the *k*_*z*_ sampling interval Δ*k*_*z*_, which is set by the real-space *z*-extent *L*_*z*_. Decreasing Δ*k*_*z*_ by increasing *L*_*z*_ improves the attainable resolution in the pitch-energy density spectrum.

## Author contributions

N.A.K. conceived the project. S.W.I. implemented the computational code. S.W.I. and J.M. curated 3D structures. J.M. visualized the results. N.M. formalized the mathematical definitions and derived theoretical results. S.W.I. and N.A.K. wrote the paper with contributions of all authors.

## Data availability

All data in this study were generated by running our codes. The structural data of DNAs and proteins were obtained from the Protein Data Bank and are available under accession codes. The structural data for ideal chiral and achiral structures will be provided as supplementary data upon acceptance. The tomographic reconstruction of nanoparticles and assemblies are available from the corresponding author upon reasonable request.

## Code availability

The code for the Fourier helicity spectra will be deposited on a designated repository and available via GitHub after acceptance.

